# Quorum sensing-dependent invasion of *Ralstonia solanacearum* into *Fusarium oxysporum* chlamydospores

**DOI:** 10.1101/2022.11.03.515128

**Authors:** Chiaki Tsumori, Shoma Matsuo, Yuta Murai, Kenji Kai

**Author notes:** These authors contributed equally to this work (Chiaki Tsumori, Kenji Kai). **Corresponding Author**, Kenji Kai.

## Abstract

Strains of *Ralstonia solanacearum* species complex (RSSC), though known as the causative agent of bacterial wilt disease in plants, induce the chlamydospores of many fungi species and invade them through the spores. The lipopeptide ralstonins are the chlamydospore inducers produced by RSSC and are essential for this invasion. However, no mechanistic investigation of this interaction has been conducted. In this study, we report that quorum sensing (QS), which is bacterial cell–cell communication, is important for RSSC to invade the fungus *Fusarium oxysporum* (*Fo*). Δ*phcB*, a deletion mutant of QS signal synthase, lost the ability to both produce ralstonins and invade *Fo* chlamydospores. The QS signal methyl 3-hydroxymyristate rescued these disabilities. In contrast, exogenous ralstonin A, while inducing *Fo* chlamydospores, failed to rescue the invasive ability. Gene-deletion and -complementation experiments revealed that the QS-dependent production of extracellular polysaccharide (EPS I) is essential for this invasion. The RSSC cells adhered to *Fo* hyphae and formed biofilms there before inducing chlamydospores. This biofilm formation was not observed in the EPS I- or the ralstonin-deficient mutant. Microscopic analysis showed that RSSC infection resulted in the death of *Fo* chlamydospores. Altogether, we reported that the RSSC QS system is important for this lethal endoparasitism. Among the factors regulated by the QS system, ralstonins, EPS I, and biofilm are important parasitic factors.

**Significance:** RSSC strains are Gram-negative bacteria that infect both plants and fungi. The *phc* QS system of RSSC is important for parasitism in plants because it allows them to invade and increase within the host by causing appropriate system activation at each infection step. In this study, we confirmed not only the importance of ralstonins as *Fo* chlamydospore inducers, but also that of biofilm formation on the hyphae. In addition to ralstonins, EPS I turned out to be important for biofilm formation. The QS system comprehensively controls the production of these factors in the interaction with *Fo*. Due to RSSC infection, the cell membranes and organelles of *Fo* chlamydospores were destroyed, showing that RSSC cells are not endosymbionts but lethal invaders. This result advocates a new QS-dependent mechanism for the process by which a bacterium invades a fungus.

## Introduction

The interactions between bacteria and fungi play a vital role in natural and agricultural ecosystems (1–3). They are cornerstone members of communities riding biochemical cycles in plant rhizosphere and contribute to plants’ health and diseases (4–6). However, our knowledge of interactions among microbes is less than that of interactions between plants and microbes. The most intimate examples of microbial interactions are bacterial parasitism/symbiosis in fungal hosts (3, 7, 8). Recent advances have shown that some bacteria can live inside fungal cells (e.g., hyphae and asexual spores) by forming endosymbiotic relationships with their host fungi. These bacteria, called endobacteria or endofungal bacteria, have now gained much attention in the studies of microbial ecology and the evolution of endocellular symbiosis. Unfortunately, there are several examples of endobacteria that cannot grow outside the fungal cells. Even when cultured, reinfection experiments have been accomplished in a limited number of cases (3, 7, 8).

The association of *Mycetohabitans rhizoxinica* (formerly *Burkholderia rhizoxinica*) with *Rhizopus* fungi is a well-studied example of endobacteria–fungus interactions (9). *Rhizopus microsporus*, a mucoromycete fungus, is the causative agent of rice seedling blight. It produces the phytotoxin rhizoxins responsible for the characteristic symptoms (10, 11). Partida-Martinez and Hertweck showed that the toxins are produced by the combination of *M. rhizoxinica* and *R. microsporus* (12, 13). The bacterial cells invade the fungal hyphae by degrading the cell walls with chitinase and may escape fungal defense using an unusual galactofuranose lipopolysaccharide (14, 15). The endobacterium-free *R. microsporus* cannot produce rhizoxins and cannot form asexual spores (12, 16). Later, the bacterial endosymbiont was found also to influence the sexual reproduction of the fungus host (17). Generally, the endobacterium is essential for the host lifestyle.

Strains of *Ralstonia solanacearum* species complex (RSSC) are known as the causative agents of bacterial wilt disease on solanaceous plants (18, 19). Studies on RSSC–host plant interactions have been accumulated. Therefore, we can report, in detail, how the bacteria invade host plant tissues and cause wilting symptoms (19, 20). Quorum sensing (QS), a bacterial cell–cell communication, plays a crucial role in RSSC virulence expression in plants, where the *phc* regulatory elements encoded by the *phcBSRQ* operon and *phcA* gene play pivotal roles (18, 19). Previously, we reported that the RSSC strain OE1-1 employs (*R*)-methyl 3-hydroxymyristate (3-OH MAME) as the *phc* QS signal (Fig. 1A) (21). The methyltransferase PhcB synthesizes 3-OH MAME, and the two-component system PhcS–PhcRQ senses the QS signal, which leads to the activation of the global virulence regulator PhcA and the subsequent *phc* QS circuit (21, 22). Deleting *phcB* or *phcA* dramatically decreased virulence in host plants (18, 22). Cell–cell chemical communication is essential for RSSC virulence in plants.

**Figure 1.**
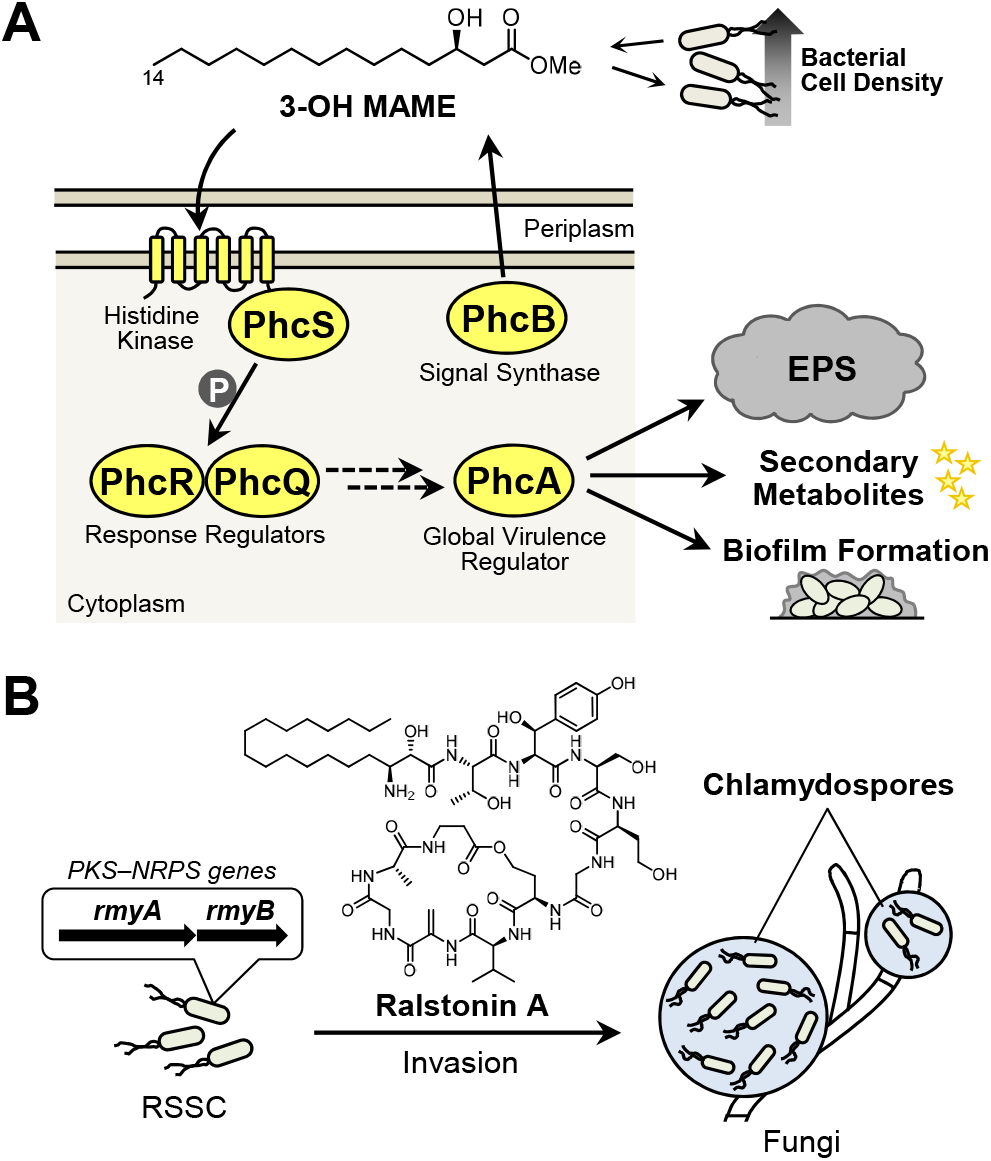
The *phc* QS system of RSSC and its endoparasitism in fungi. (A) Schematic diagram of the *phc* QS system in strain OE1-1. As bacterial density increases, 3-OH MAME concentrations rise, and the molecule is received by PhcS–PhcRQ. Then, PhcA is functionalized and induces the production of virulence factors, such as EPS, secondary metabolites, and biofilms. (B) The possible endoparasitism of RSSC coupled with chlamydospore induction. Ralstonin A synthesized by the PKS–NRPS induces chlamydospore formation in many fungi and may facilitate endoparasitism.

In 2016, Keller and co-workers found that RSSC strain GMI1000 can induce chlamydospore formation in phylogenetically diverse fungi (including members of Mucoromycota, Ascomycota, and Basidiomycota) and invade the fungal cells probably through the formed chlamydospores (Fig. 1B) (23). They also suggested that the polyketide synthase–nonribosomal peptide synthetase (PKS–NRPS) enzymes encoded by *rmyA*/*rmyB* genes may produce a pivotal compound in inducing chlamydospores. We later identified ralstonins A and B, unique cyclic lipopeptides from RSSC strains OE1-1 and GMI1000, to be the chlamydospore inducers for *Fusarium oxysporum* (Fig. 1B) (24). One month later, Prof. Nett and co-workers also reported the chemical characterization of the ralstonins (aka ralsolamycins) (25). However, because we have just began facing the results of these RSSC–fungi interactions, little is still known on how and why the bacteria invade the fungal cells. Is there some commonality between this endoparasitism and plant-wilt disease development? Is the *phc* QS system of RSSC involved in endoparasitism?

A quantitative assay for evaluating the invasion rate of RSSC cells into *F. oxysporum* chlamydospores was developed in this study. We investigated the key factors establishing this interkingdom interaction by analyzing the invasion rates of the gene-deletion and -complementation RSSC mutants. A coculturing assay to directly observe the interaction between RSSC and *F. oxysporum* was extablished. Consequently, ralstonin/EPS I production and biofilm formation, both of which are regulated by the *phc* QS system, turned out to be essential. Alternatively, extracellular enzymes and type II/III secretion systems, although these are important infecting plants, are not involved. An insight into the behaviors of RSSC cells after the invasion of *F. oxysporum* chlamydospores was obtained using TEM (transmission electron microscope) and fluorescence microscope. Based on these findings, snapshots of the course of the QS-dependent entry of RSSC cells into *F. oxysporum* chlamydospores was proposed.

## Results

### Ralstonins are required for inducting *F. oxysporum* chlamydospores and the RSSC’s invasion into them

RSSC strain OE1-1 aggressively induces the chlamydospore formation of *F. oxysporum* NBRC 31213 and invades them when these microorganisms are cocultured (24, 26). *F. oxysporum* strains form fewer conidia than *Asperigillus* spp., making it suitable for microscopic observation. A nylon net sandwich assay that assessed the invasion rate of this bacterium into the chlamydospores of *F. oxysporum* using a constitutive GFP-producing vector was developed (pDSK-GFP) (Fig. 2A). In this method, the invasion rate can be calculated efficiently by washing the nylon net to remove excess RSSC cells and collecting the formed chlamydospores with a cover glass. Strain OE1-1 (wild-type) showed 8.8% invasion into the formed chlamydospores (Fig. 2B). The invasion rate of the wild-type strain was around 10% throughout the experiments conducted in this paper. Δ*rmyA*, a ralstonin-deficient mutant of strain OE1-1 (24, 27), did not induce the chlamydospores and thus could not establish the endoparasitism (Fig. 2B). The exogenous ralstonin A (0.1 and 1 nmol/disc) rescued the chlamydospore formation and endoparasitic invasion of Δ*rmyA*. According to these results, ralstonin-dependent chlamydospore formation is needed for RSSC’s entry into the spores and this assay can quantitatively evaluate the invasion rates of RSSC strains.

**Figure 2.**
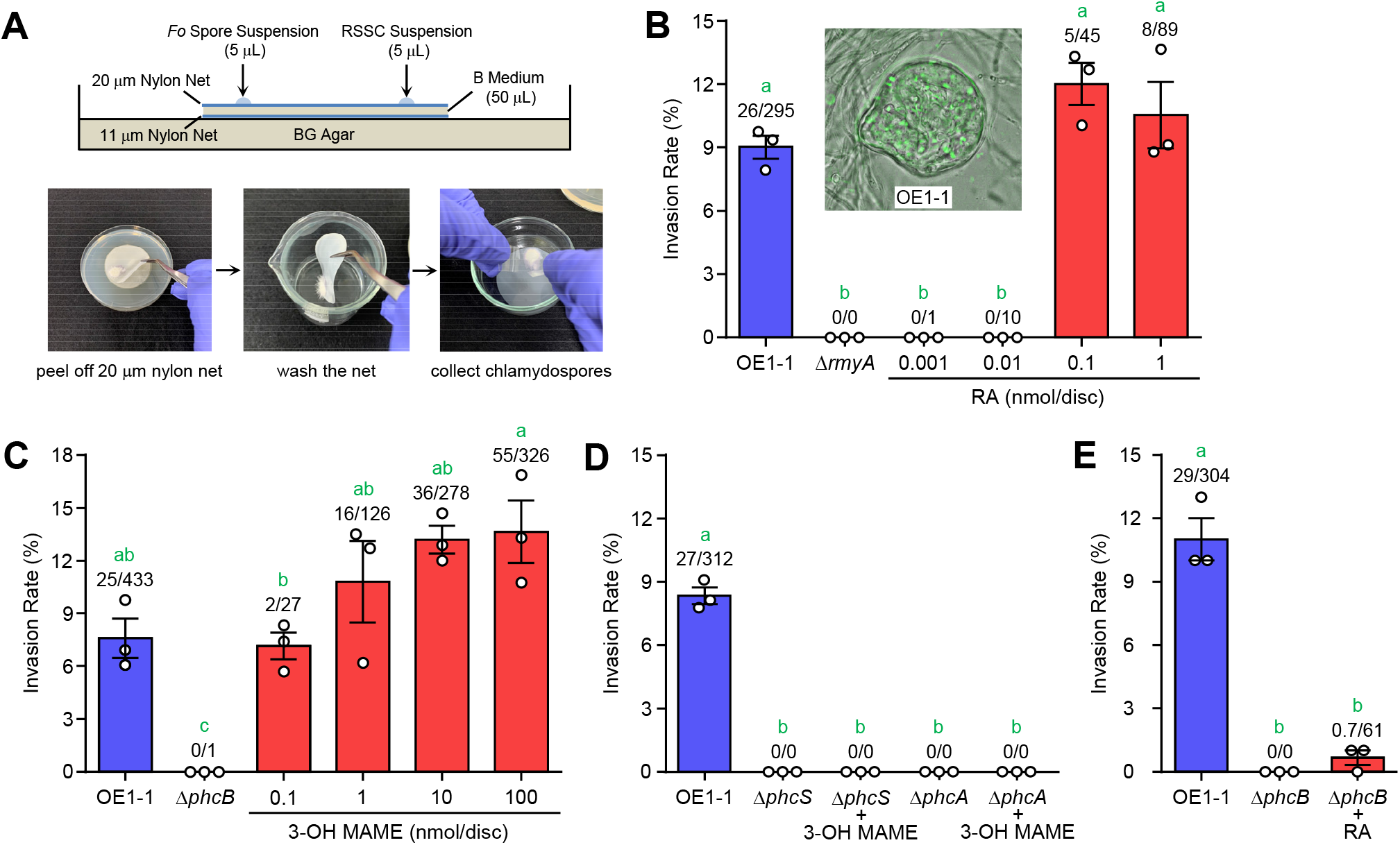
Analysis of endoparasitism of the RSSC strain OE1-1 and its mutants in *F. oxysporum*. (A) A nylon net sandwich assay to evaluate the invasion rate of RSSC strains. The images of assay plate (*upper*) and chlamydospore recovery method (*lower*) are shown. (B) Invasion rates of OE1-1 and Δ*rmyA* into *F. oxysporum* chlamydospores. The addition of ralstonin A (RA) rescued the invasion rate of Δ*rmyA*. A merged photo of a chlamydospore infected with OE1-1 is shown as the inset. (C) Evaluation of the invasion ability of Δ*phcB* into *F. oxysporum* chlamydospores and the effect of 3-OH MAME. The addition of 3-OH MAME rescued the invasion rate of Δ*phcB*. (D) Invasion rates of Δ*phcS* and Δ*phcA* and their response to 3-OH MAME (1 nmol/disc). (E) The effect of ralstonin A (0.1 nmol/ disc) on the invasion of Δ*phcB*. The error bars show the mean ± SEM (*n* = 3). The numbers above the bars indicate infected spores/total spores. The green letters above the bars show a significant difference (*p* < 0.05, Tukey’s test).

### QS-Deficient mutants loss invasion ability into *F. oxysporum* chlamydospores

The virulence of RSSC on plants is tightly regulated by the *phc* QS system (18, 22). Therefore, we wondered whether the *phc* QS system is also involved in the interaction of RSSC with *F. oxysporum*. To examine this, we investigated the invasion ability of Δ*phcB* (a QS signal synthase-deletion mutant) (21) into *F. oxysporum. ΔphcB* neither induced the chlamydospore formation of *F. oxysporum* nor established the endoparasitism on the fungus (Fig. 2C). Exogenous 3-OH MAME dose-dependently rescued the invasion rate of Δ*phcB*. When 3-OH MAME was applied at 0.1 nmol/disc, Δ*phcB* exhibited a similar invasion rate with the wild-type strain. Also, 3-OH MAME increased the invasion rate over the wild-type strain at higher concentrations. Δ*phcS* (QS signal receptor-deficient) and Δ*phcA* (the global QS regulator deficient) (21, 28) also lost the invasion ability into the chlamydospores, which was not restored by 3-OH MAME supplement (Fig. 2D). According to these results, the *phc* QS circuit signaling activation by 3-OH MAME is important for RSSC invasion into *F. oxysporum* chlamydospores.

The production of ralstonins is under the control of the *phc* QS system in RSSCs (24). Furthermore, we checked whether the loss in invasion ability of Δ*phcB* and the related mutants is attributed only to the deficiency of ralstonin production. Consequently, while ralstonin A (0.1 nmol/disc) induced the formation of many chlamydospores, the compound did not rescue the invasion rate of Δ*phcB* to that of strain OE1-1 (Fig. 2E). The concentration of ralstonin A completely rescued the invasion rate of Δ*rmyA*. The same results for Δ*phcB* were obtained for Δ*phcS* and Δ*phcA* (Fig. S1). Therefore, it was concluded that other factor(s) regulated by the *phc* QS system are also crucial for establishing the RSSC parasitism.

### Extracellular enzymes and type II/III secretion systems are not involved in RSSC invasion

To invade *F. oxysporum* chlamydospores, the RSSC cells should overcome their physical barriers (e.g., cell membranes and walls) (15). As expected, this bacterium harbors two chitinase genes (*RSp0275* and *RSp0924*), three glucanase genes (*cbh-A, egl*, and *RSc0818*), and two lipase (or esterase) genes (*RSp0138* and *RSp0161*) (29). The previous RNA-Seq comparison between OE1-1 and Δ*phcB* showed that the expression of these genes is under the control of the *phc* QS system (30–32). We thus constructed these gene-deletion mutants and evaluated their invasion ability into *F. oxysporum* chlamydospores. However, the deletion mutants of these genes invaded the chlamydospores with a similar rate as the wild-type strain (Fig. 3A). Because many extracellular enzymes (including uncharacterized ones) are secreted by type II secretion system (T2SS) (19), we created its deficient mutant (Δ*gspD; gspD* encodes an important protein of T2S machinery) (33) and conducted the invasion assay. Δ*gspD* also showed a similar invasion rate as strain OE1-1 (Fig. 3A). Fungal cell membranes and walls may not be a barrier for RSSC to accomplish this invasion, or they may be using a completely different method. We concluded that such extracellular enzymes are not critical parasitic factors regulated by the *phc* QS system.

**Figure 3.**
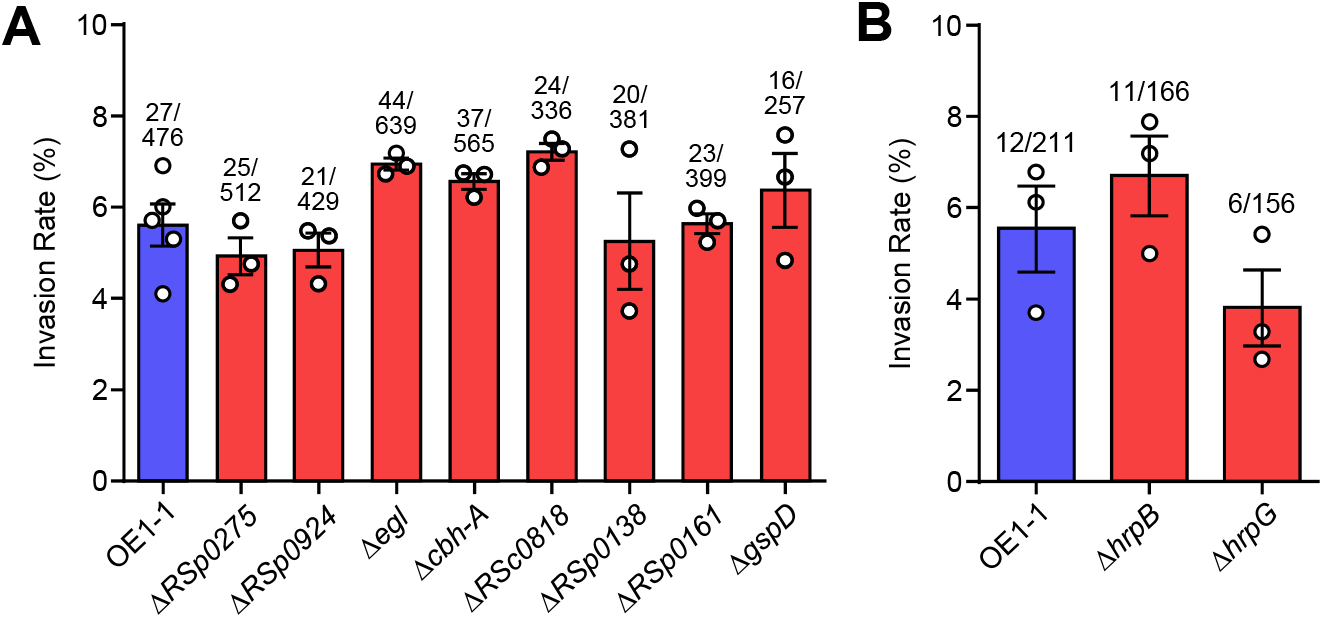
Evaluation of extracellular enzymes, T2SS, and T3SS on RSSC endoparasitism in *F. oxysporum*. (A) Invasion rates of the gene-deletion mutants of extracellular enzymes and the T2SS component into *F. oxysporum* chlamydospores. Chitinase genes (*RSp0275* and *RSp0924*), glucanase genes (*cbh-A, egl*, and *RSc0818*),lipase genes (*RSp0138* and *RSp0161*), and a T2SS machinery gene (*gspD*). (B) Invasion rates of Δ*hrpB* and Δ*hrpG* into *F. oxysporum* chlamydospores. The error bars show the mean ± SEM (*n* = 3 or 5). The numbers above the bars show infected spores/total spores. There were no statistically significant differences in the values of each data set (*p* < 0.05, Tukey’s test).

Previously, the importance of type III secretion system (T3SS) in *M. rhizoxinica–R. microsporus* interaction was described (34). T3SS is a specialized syringe-shaped protein-export system used by several pathogenic Gram-negative bacteria to inject virulence proteins (effectors) into host cells and in many cases, it is important for bacterial pathogenicity (35, 36). In RSSC, the expression of T3SS components is regulated by the *phc* QS system and is up-regulated or down-regulated depending on the culturing conditions (37, 38). We examined the invasion rates of Δ*hrpB* and Δ*hrpG* to check the importance of T3SS in RSSC–*F. oxysporum* interaction. HrpB and HrpG are the positive regulators of the RSSC T3SS, and HrpG is the higher priority (39, 40). Δ*hrpB* did not show a significant decrease in invasion rates into *F. oxysporum* chlamydospores compared to the wild-type strain, while Δ*hrpG* still showed invasion the ability of the fungus (Fig. 3B). This showed that RSSC does not use T3SS for this invasion and that a factor regulated by HrpG may have a small effect on the invasion rate. Altogether, the mechanism of invasion of RSSCs into *F. oxysporum* was shown to be different from that of *M. rhizoxinica*–*R. microsporus* interaction. The endoparasitic factors controlled by the *phc* QS system remain a mystery.

### EPS I is important for RSSC invasion into *F. oxysporum* chlamydospores

The production of EPS I (a major extracellular polysaccharide of RSSC) and LecM (an RS-IIL lectin) is also dependent on the activation of the *phc* QS system (31, 32, 37). As shown in *Pseudomonas aeruginosa* (41), EPS and lectin may comprise the extracellular matrix of RSSC cells. To investigate the importance of EPS I and LecM, we examined the invasion rates of Δ*epsB* (EPS I deficient) and Δ*lecM* (LecM deficient). These defective mutants were not parasitic on *F. oxysporum*, whereas the gene-complemented strains (*epsB*-comp and *lecM*-comp) were (Fig. 4A). We previously confirmed that the *phc* QS system in Δ*epsB* and Δ*lecM* does not drive appropriately due to an unveiled feedback regulation (42, 43). To investigate the true importance of EPS I and LecM, we complemented the production of these factors in Δ*phcB* (named them Δ*phcB-CxpsR* and Δ*phcB-ClecM*, respectively) and examined the invasion rates of these gene-complimented strains. Because XpsR is a transcriptional regulator for *eps* biosynthetic gene cluster in RSSC (44, 45), Δ*phcB-CxpsR* constitutively produces EPS I (Fig. 4B). The result of invasion assays showed that Δ*phcB-CxpsR* showed higher invasion rate than strain OE1-1 in the presence of ralstonin A, though Δ*phcB* and Δ*phcB-ClecM* could not invade the chlamydospores (Fig. 4C). In conclusion, EPS I production only is vital for RSSC entry into *F. oxysporum* chlamydospores.

**Figure 4.**
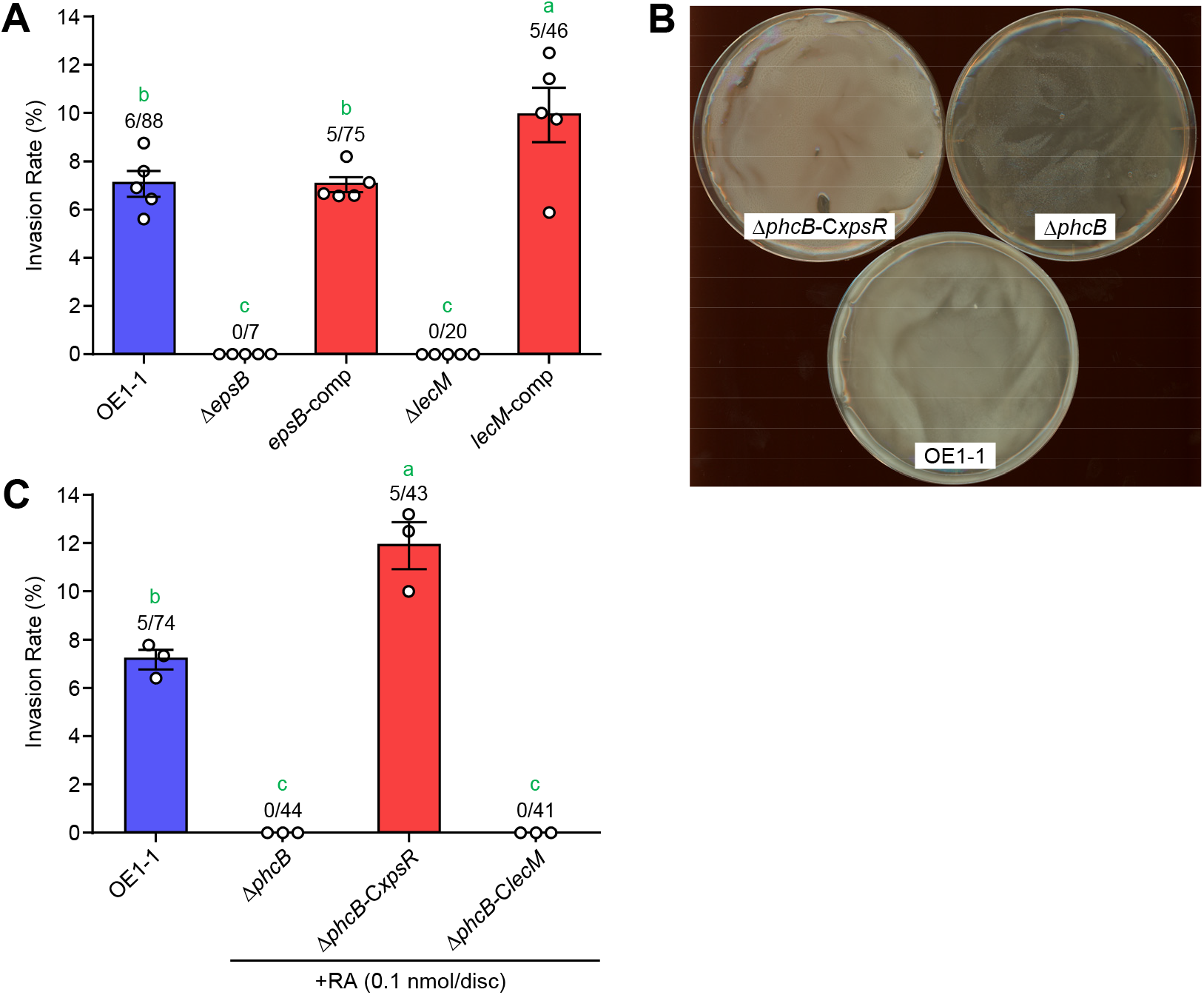
Effects of EPS I and LecM on the RSSC endoparasitism in *F. oxysporum*. (A) Invasion rates of Δ*epsB, epsB*-comp, Δ*lecM*, and *lecM*-comp into *F. oxysporum* chlamydospores. (B) The colonies of Δ*phcB-CxpsR*, Δ*phcB*, and OE1-1 on BG agar plates. The number of EPS I produced differed among the strains (Δ*phcB* < OE1-1 < Δ*phcB-CxpsR*). (C) Invasion rates of Δ*phcB*, Δ*phcB-CxpsR*, and Δ*phcB-ClecM* into *F. oxysporum* chlamydospores. Ralstonin A (RA) was used to induce *F. oxysporum* chlamydospores. The error bars show the mean ± SEM (*n* = 3 or 5). The numbers above the bars show infected spores/total spores. The green letters above the bars show significant differences (*p* < 0.05, Tukey’s test).

### EPS I is important for the biofilm formation of RSSC cells on *F. oxysporum* hyphae

To investigate how EPS I is involved in the RSSC parasitism, we observed the interaction between OE1-1 and *F. oxysporum* during a coculture assay in a glass-bottom dish (Fig. 5A) and looked for event(s) that EPS I might be involved in. We found that RSSC cells formed biofilm-like structures on the fungal hyphae before inducing the chlamydospores (Fig. 5B). This implied that the bacterial biofilms on the host may be involved in RSSC–*F. oxysporum* interaction. The QS-deficient mutants, Δ*phcA* and Δ*phcB*, did not adhere to the mycelia of *F. oxysporum*, nor did they form biofilms there (Fig. 5C). Thus, the cell attachment and biofilm formation, regulated by the *phc* QS system (22, 38), may be essential traits for the RSSC parasitism on the host hyphae.

**Figure 5.**
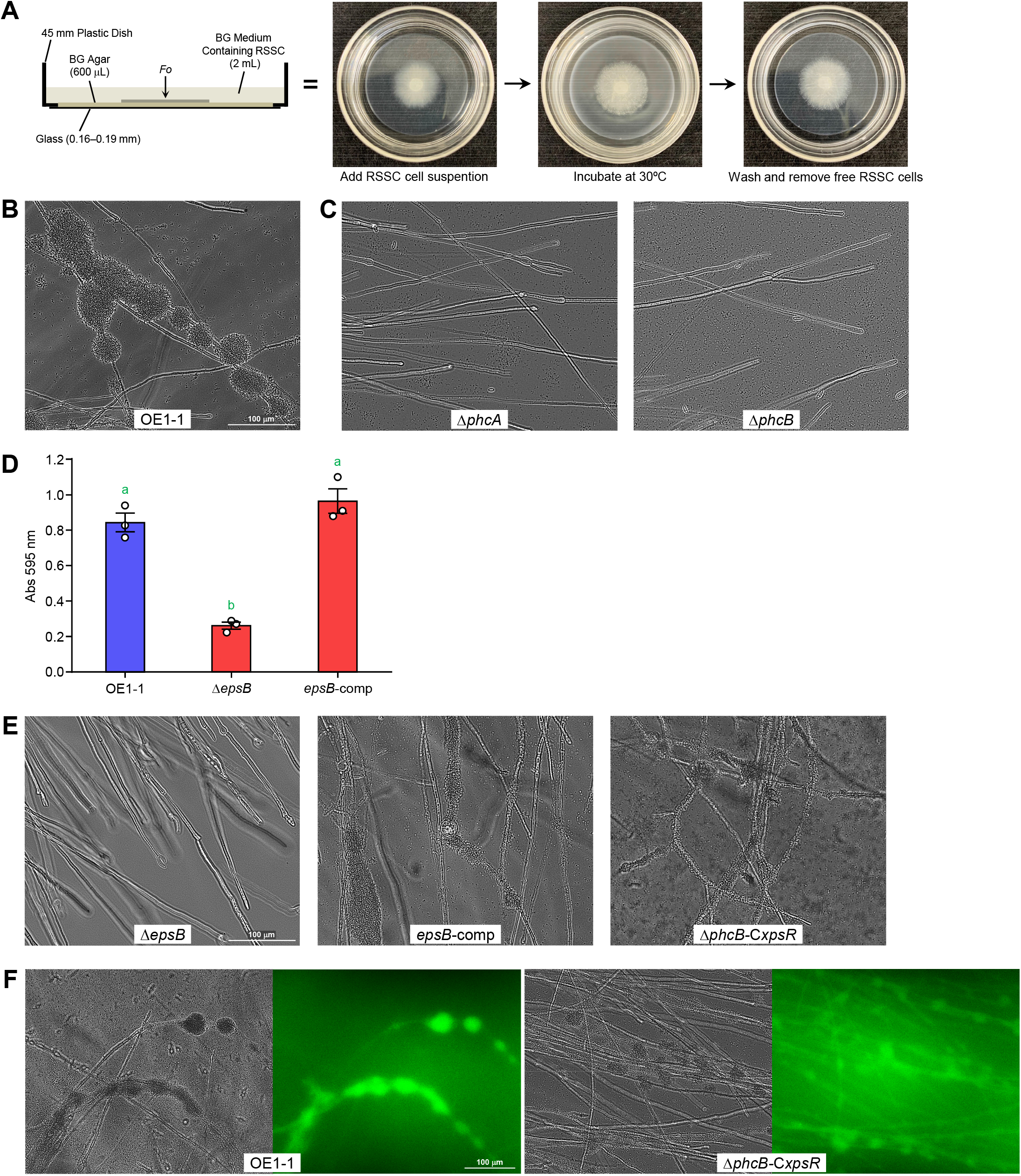
Importance of EPS I on RSSC biofilms’ formation on *F. oxysporum* hyphae.(A) A glass-bottom dish assay to observe the interaction between RSSC and *F. oxysporum*. The images of the assay plate (*left*) and the assay method (*right*) are shown. (B) The biofilms of OE1-1 on *F. oxysporum* hyphae. RSSC cells form biofilms like a grape cluster on the hyphae. (C) The photos of coculturing of *F. oxysporum* with Δ*phcA* and Δ*phcB*. These mutants did not form biofilms. (D) The biofilms of OE1-1, Δ*epsB*, and *epsB*-comp on a plastic plate. The vertical axis represents the amount of biofilm as the absorbance of the recovered crystal violet. The error bars show the mean ± SEM (*n* = 3). The green letters above the bars shows significant differences (*p* < 0.05, Tukey’s test). (E) The photos of coculturing of *F. oxysporum* with Δ*epsB, epsB*-comp, and Δ*phcB-CxpsR*. The biofilm formation was observed in *epsB*-comp and Δ*phcB-CxpsR*.(F) DBA-FITC staining of the biofilms of OE1-1 and Δ*phcB-CxpsR*. The bright field image is on the left, and the fluorescence image is on the right.

The attachment of RSSC cells to solid surfaces (such as plastic) and the formation of biofilms there are dependent on the production of EPS I (Fig. 5D) (46). However, there was no knowledge of whether EPS I is involved in the RSSC adhesion to fungal mycelia and biofilm formation. The influence of *epsB* deletion or complementation was evaluated by coculture assays with *F. oxysporum*. Although Δ*epsB* did not form the biofilm on *F. oxysporum* hyphae, it did form the biofilm after the complementation of *epsB* gene (Fig. 5E). Notably, Δ*phcB-CxpsR* showed the ability to form biofilms at a high frequency on *F. oxysporum* hyphae, although the biofilm was thinner than that of the wild-type strain. Therefore, we expected that even with increased EPS I levels, the biofilms would not mature due to the absence of other factors (such as LecM) in Δ*phcB*-C*xpsR*. Additionally, when the fluorescent staining of EPS I was conducted using DBA-FITC (*Dolichos biflorus* agglutinin, a lectin with specificity for α-linked *N*-acetylgalactosamine) (47), The FITC-derived fluorescence was observed from the biofilms of OE1-1 and Δ*phcB-CxpsR* (Fig. 5F). Due to the increased staining and washing operations, the biofilms are somewhat more broken than those in Figure 5B and E. This showed that the bacteria produced EPS I there to form biofilms. EPS I production was definitely found to be essential for the biofilm formation of RSSC cells on *F. oxysporum* hyphae.

### Quality and quantity of EPS I are essential for RSSC endoparasitism

We predicted that the EPS I-dependent biofilm formation of RSSC cells may be necessary for their parasitism on *F. oxysporum*. To examine this, we tested the parasitic potential of mutant strains lacking the enzymes involved in EPS I biosynthesis, that is, to change the substructure of EPS I (Fig. 6A). RSp1012, a LpxA-like protein, transfers a 3-hydroxybutanoate to the 2,4-diamino-2,4,6-trideoxygalactose moiety in EPS I (48). Δ*RSp1012* produced the EPS I that differed in appearance from that of the wild-type strain (Fig. 6B). The mutant formed a more brittle biofilm than OE1-1 on *F. oxysporum* hyphae, and their thickness became thinner (Fig. 6C, *upper*). Δ*RSp1012* parasitized the fungus at almost the same rate as the wild-type strain (Fig. 6 D). The deletion of the gene *RSp1007* encoding the enzyme that transfers the acetate to amino sugars (Fig. 6A) (49) resulted in reduced EPS production and no invasion of the mutant into *F. oxysporum* chlamydospores (Fig. 6B and D). Ralstonin production was reduced in Δ*RSp1007* (Fig. S2). This may be obtained from the feedback inhibition observed in Δ*epsB* and Δ*lecM* (42, 43). Thus, the invasion assay of Δ*RSp1007* with ralstonin A was conducted. A certain recovery of the parasitism rate was observed, but not to the level of the wild-type strain (Fig. 6E). In a glass-bottom dish assay, Δ*RSp1007* formed a fragile biofilm on *F. oxysporum* mycelium, and the frequency of its formation decreased slightly (Fig. 6C, *lower*). In a biofilm test using a plastic assay plate, both mutants formed slightly more biofilms than Δ*phcB*, and the amounts were greatly reduced compared to OE1-1. (Fig. 6F). The biofilm on plastic was probably more influenced by the quality and quantity of EPS I. We showed that the EPS I, which acts as an adhesive to some extent, is essential for RSSC to parasitize *F. oxysporum* and to drive the QS circuit appropriately.

**Figure 6.**
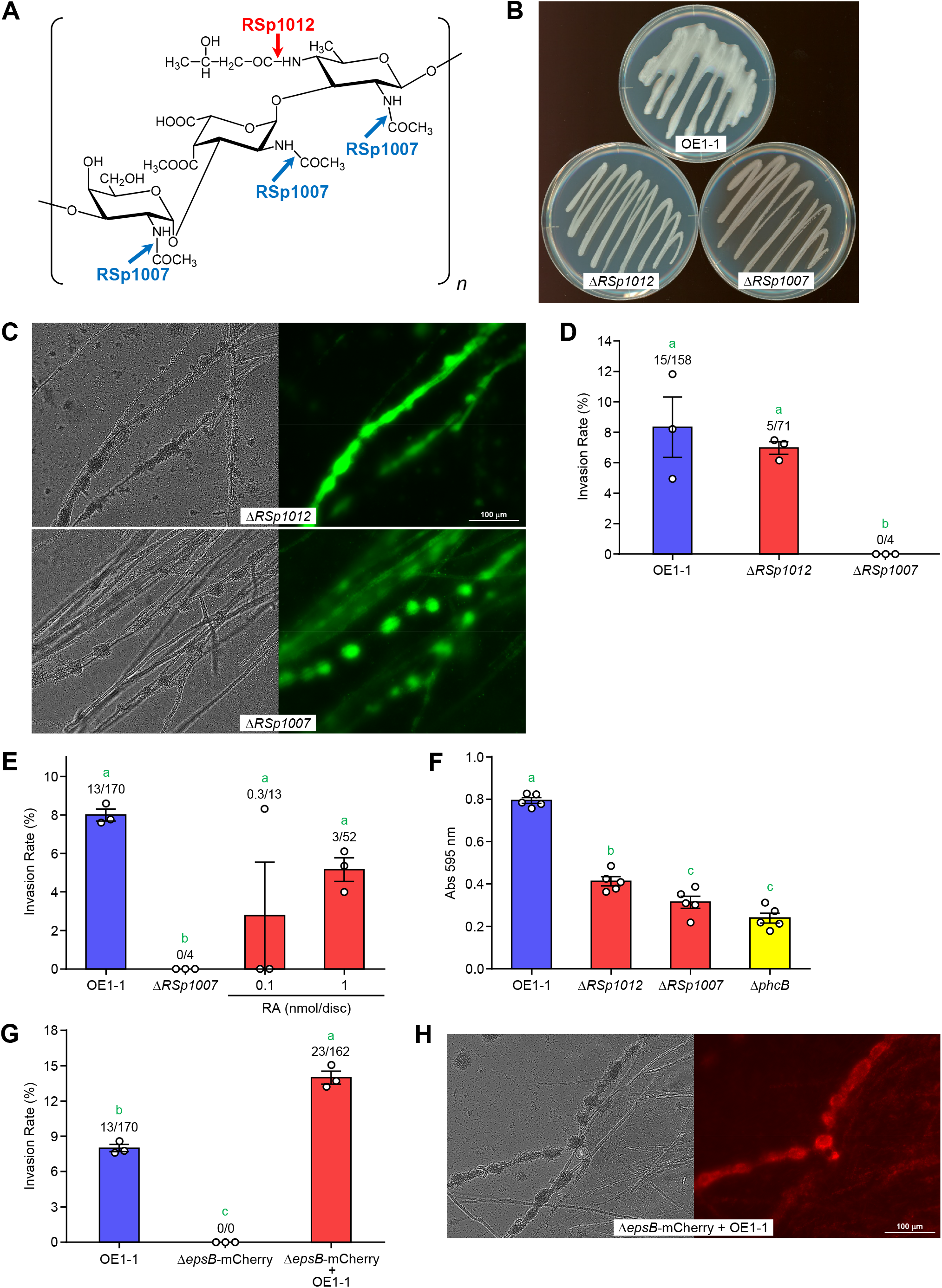
Importance of EPS I structure on RSSC endoparasitism in *F. oxysporum* and recovery of Δ*epsB* endoparasitism ability by coculturing with OE1-1. (A) The structure of EPS I is produced by RSSC strains. RSp1012 and RSp1007 are involved in transfer reactions at the positions shown by the arrows. (B) A photo image of OE1-1, Δ*RSp1012*, and Δ*RSp1007* grown on BG agar plates. Colony liquidity derived from EPS I is significantly reduced in Δ*RSp1012* and Δ*RSp1007*. (C) The biofilms of Δ*RSp1012* and Δ*RSp1007* on *F. oxysporum* hyphae. SYTO9 was used to stain bacterial cells. The bright field image is on the left, and the fluorescence image is on the right. (D) Invasion rates of OE1-1, Δ*RSp1012*, and Δ*RSp1007* into *F. oxysporum* chlamydospores. (E) The effect of ralstonin A (RA) on the invasion of Δ*RSp1012* into *F. oxysporum* chlamydospores. (F) The biofilms of OE1-1, Δ*RSp1012*, Δ*RSp1007*, and Δ*phcB* were on a plastic plate. The vertical axis represents the amount of biofilm as the absorbance of the recovered crystal violet. (G) Invasion rates of OE1-1, Δ*eps*-mCherry, and Δ*eps*-mCherry cocultured with OE1-1 into *F. oxysporum* chlamydospores. (H) The biofilm formation by OE1-1 and Δ*eps*-mCherry on *F. oxysporum* hyphae. The bright field image is on the left, and the fluorescent image is on the right. The error bars show the mean ± SEM (*n* = 3 or 5). The numbers above the bars show infected spores/total spores. The green letters above the bars show significant differences (*p* < 0.05, Tukey’s test).

To determine when EPS I was significant, we tried to mix Δ*epsB* and OE1-1 and to examine the parasitism of Δ*epsB* on *F. oxysporum*. To distinguish between Δ*epsB* and wild-type cells, mCherry protein was expressed in Δ*epsB* (named Δ*epsB*-mCherry). mCherry-derived fluorescence was observed in the *F. oxysporum* chlamydospores from coculturing experiments, confirming that Δ*epsB*-mCherry is taken up by the spores (Fig. 6G). Furthermore, the invasion rate was higher than that of the wild-type strain. Next, we examined whether Δ*epsB* was incorporated into the biofilms of OE1-1 during a coculture assay. Δ*epsB*-mCherry along with the wild-type strain, formed mature biofilms on *F. oxysporum* hyphae, although the uptake ratio of the mutant varied from biofilm to biofilm (Fig. 6H). When *E. coli* DH5α-mCherry and OE1-1 were cocultured, the *E. coli* strain was not incorporated into biofilms or the *F. oxysporum* chlamydospores (Fig. S3). This result suggested that not all types of bacteria can be incorporated into the biofilm of RSSC and can invade *F. oxysporum* chlamydospores. Altogether, it was exhibited that EPS I-dependent biofilm formation is important for RSSC invasion, but EPS I may not be essential for the further invasion process.

### Ralstonin is also involved in the biofilm formation of RSSC on *F. oxysporum* hyphae

According to reports, lipopeptides are involved in the biofilm formation of Gram-negative bacteria as biosurfactants (50, 51). Therefore, we investigated whether ralstonins are responsible for the biofilm formation in RSSC cells. Δ*rmyA* did not form biofilms on the hyphae of *F. oxysporum*, and the biofilm formation was partially restored with the addition of ralstonin A (0.1 and 1 μM) (Fig. 7A). However, no biofilm was observed at 10 μM, suggesting that high concentrations of ralstonin A may act rather inhibitory. Instead, chlamydospores were formed faster due to increased ralstonin A concentration. Regarding the biofilm formation assay using a plastic assay plate, Δ*rmyA* resulted in less biofilm than the wild-type strain, and ralstonin addition rescued the biofilm formation of Δ*rmyA* (Fig. 7B). Therefore, ralstonins play an essential role in the induction of *F. oxysporum* chlamydospore formation and in the RSSC’s biofilm formation on *F. oxysporum* mycelia.

**Figure 7.**
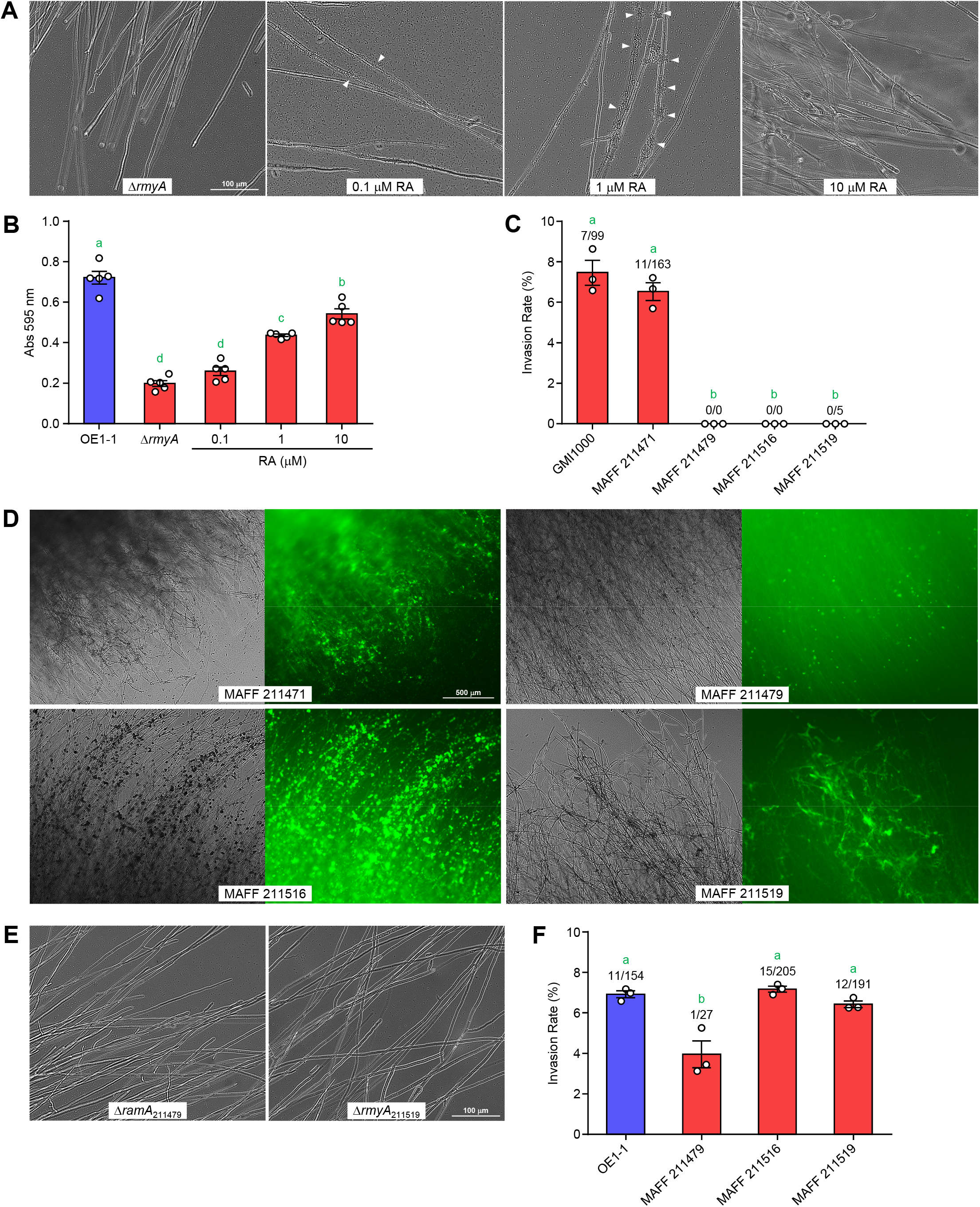
The importance of lipopeptides on RSSC endoparasitism in *F. oxysporum*. (A) Ralstonin-dependent recovery of biofilm formation in Δ*rmyA*. Ralstonin A (RA) supplement at 10 μM caused the chlamydospore formation of *F. oxysporum*. (B) Biofilm formation of Δ*rmyA* on a plastic plate and its response to ralstonin A. The vertical axis represents the amount of biofilm as the absorbance of the recovered crystal violet. (C) Invasion rates of the wild-type RSSC strains into *F. oxysporum* chlamydospores. GMI1000 and MAFF 211471 produce ralstonins, MAFF 211479 and MAFF 211516 produce ralstoamides, and MAFF 211519 produces ralstopeptins. (D) The biofilms of MAFF strains on *F. oxysporum* hyphae. SYTO9 was used to stain the bacterial cells. The bright field image is on the left, and the fluorescent image is on the right. (E) Lack of biofilm formation ability in Δ*ramA* and Δ*rmyA*_211519_ on *F. oxysporum* hyphae. (F) Acquisition of endoparasitic ability of MAFF strains by coculturing with OE1-1. Only OE1-1is a positive control. The error bars show the mean ± SEM (*n* = 3). The numbers above the bars show infected spores/total spores. The green letters above the bars show significant differences (*p* < 0.05, Tukey’s test).

Some RSSC strains produce ralstopeptins or ralstoamides, which have different structures from ralstonins (Fig. S4) (52, 53). RSSC lipopeptides other than ralstonins did not show strong chlamydospore-inducing activity (53). Are they involved in biofilm formation on *F. oxysporum* mycelia? To investigate this, other RSSC strains producing ralstonins, ralstoamides, or ralstopeptins were cocultured with *F. oxysporum*. MAFF 211471 and GMI1000, which produce ralstonins, exhibited an invasion rate close to that of OE1-1, but ralstoamide-producing strains (MAFF 211479 and MAFF 211516) and ralstopeptin-producing strain (MAFF 211519) failed to establish the endoparasitism (Fig. 7C). This may be because little or no chlamydospore formation occurred in the assays with MAFF 211479, MAFF 211516, and MAFF 211519. These MAFF strains formed biofilms on the mycelia in a glass-bottom dish assay, although their ability to do so differed (Fig. 7D). The biofilms were easily collapsed by the washing process compared with strain OE1-1. Δ*rmyA*_211519_ (a ralstopeptin-deficient mutant of MAFF 211519) (53) and Δ*ramA*_211479_ (a ralstoamide-deficient mutant of MAFF 211479) (52) were found to have lost the ability to form biofilms (Fig. 7E). When these strains were tested together with OE1-1 for parasitism into *F. oxysporum*, the MAFF 211516 and MAFF 211519 exhibited invasion rates comparable to that of the OE1-1 strain (Fig. 7F). MAFF 211479 appeared to exhibit low invasion ability, even if OE1-1 was present. Overall, the lipopeptides produced by RSSC strains were shown to be essential for biofilm formation on *F. oxysporum* mycelia. Again, ralstonins were found to be essential for this endoparasitism.

### RSSC infection kills *F. oxysporum* chlamydospores

It was expected that TEM observation of chlamydospores infected by RSSC may provide some insight into the invasion mechanism and the post-invasion behaviors of RSSC. The TEM observation of chlamydospores induced by ralstonin A showed that the structures were almost identical to those previously reported (Fig. 8A) (53, 54). Particularly, the cell wall was thick, and large lipid droplets, nucleus, and ribosomes were observed in the spores. The spores were histologically confirmed to be *F. oxysporum* chlamydospores. Next, *F. oxysporum* chlamydospores infected with RSSC were observed by TEM. TEM observation of the chlamydospores after three days of cocultivation showed that the spores had disrupted cell membranes, which changed to membrane vesicles (Fig. 8B). Furthermore, organelles such as lipid droplets and nucleus disappeared, and some to many RSSC cells were observed in the chlamydospores. Bigger polyhydroxyalkanoate granules (55, 56) were observed in the RSSC cells, suggesting that the chlamydospore contents may be used as nutrients. These observations showed that these chlamydospores were killed by the RSSC’s infection and growth.

**Figure 8.**
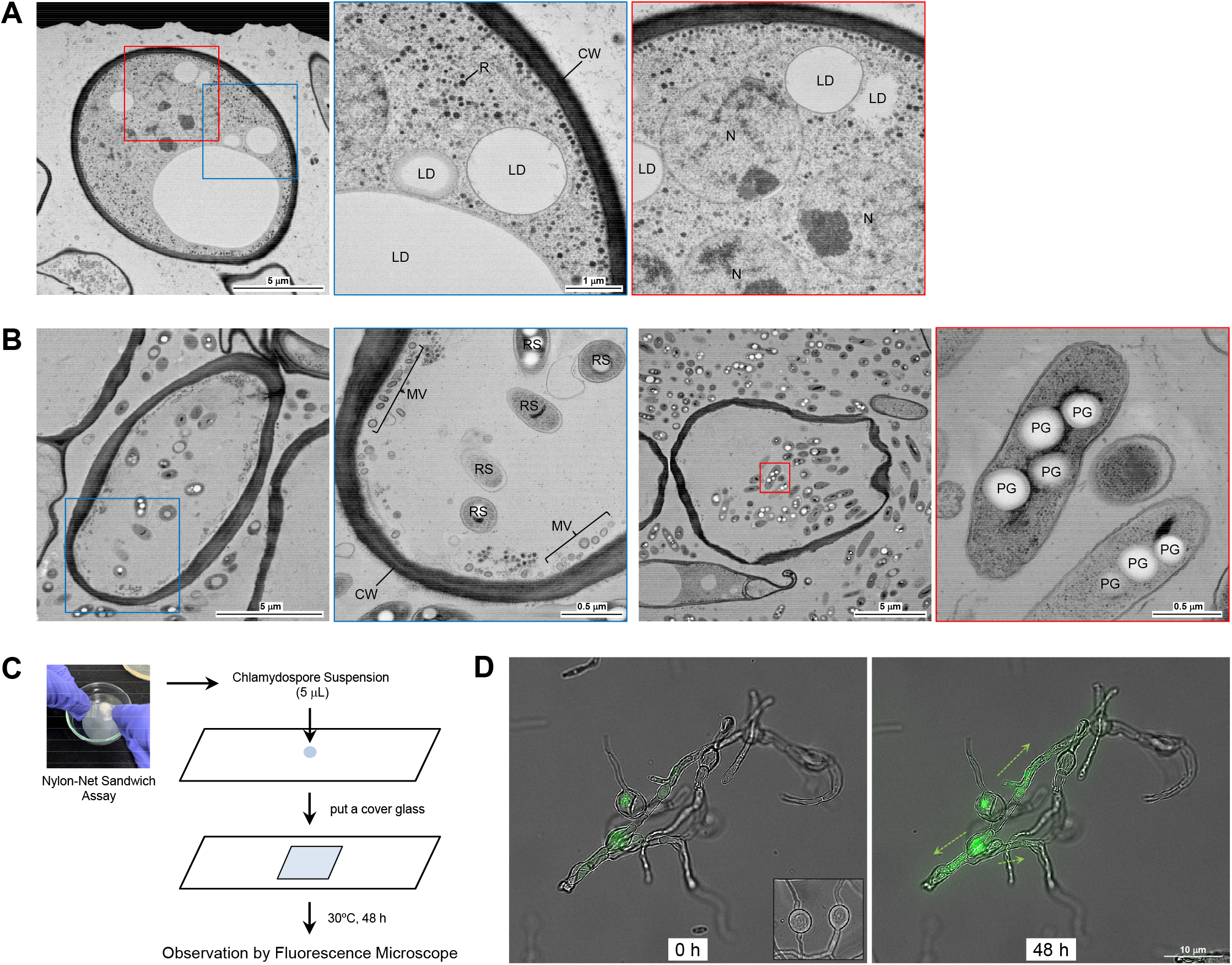
Microscopic observation of intact and infected *F. oxysporum* chlamydospores. (A) TEM images of *F. oxysporum* chlamydospores induced by ralstonin A treatment. Enlarged views of the areas framed in blue and red are shown on the right. N: nuclear, LD: lipid droplet, R: liposome, CW: cell wall. (B) TEM images of *F. oxysporum* chlamydospores infected with OE1-1. Enlarged views of the areas framed in blue or red are shown on the right. RS: RSSC cell, MV: membrane vesicle, PG: polyhydroxyalkanoate granule. (C) Preparation of a sample for the observation of RSSC behavior in *F. oxysporum* chlamydospores. (D) Fluorescence observation of infected chlamydospores immediately after collection and 24 h later. Inset shows the chlamydospores induced by ralstonin A.

We examined whether the cell death of *F. oxysporum* chlamydospores, as observed in the TEM analysis, is also observed in the unfixed chlamydospores after RSSC infection. We used a method that allowed us to observe the parasitized spores under high magnification by a fluorescence microscope (Fig. 8C). The *F. oxysporum* spores infected with OE1-1 were collected and allowed to grow on a sliding glass. The OE1-1-derived GFP fluorescence was observed over time. The RSSC cells were found to grow within infected chlamydospores and migrate into nearby hyphae (Fig. 8D). Notetably, what appeared to be cell membranes within the chlamydospores deviated from the cell walls. Alternatively, the chlamydospores induced by ralstonin A were almost spherical and showed no signs of separation of the cell membranes from the cell walls (Fig. 8D, *inset*). Therefore, we concluded that the chlamydospores infected by RSSC cells were dead, and the bacteria could migrate to the surrounding mycelia.

## Discussion

This study demonstrates that the *phc* QS system is important for RSSC to invade *F. oxysporum*. Δ*phcB* lost both the abilities to produce ralstonins and to invade *F. oxysporum* chlamydospores, and these defects were restored by the exogenous 3-OH MAME but not by ralstonin A. Analysis of the gene-deletion and -complemented mutants showed that, in addition to ralstonins, EPS I is an important parasitic factor. Moreover, EPS I and ralstonins are required for RSSC biofilm formation on *F. oxysporum* hyphae. Biofilm formation is essential for RSSC endoparasitism in *F. oxysporum*.

Based on these results, we proposed a snapshot mechanism by which RSSC cells parasitize into *F. oxysporum* via the chlamydospores induced (Fig. 9). RSSC cells exhibit positive chemotaxis toward *F. oxysporum* hyphae and migrate toward them. This can be seen in the post coculture photo in Figure 5A. After adherence, RSSC cells begin to drive the *phc* QS system. Consequently, they started to produce EPS I and ralstonins and, simutaneously, form a biofilm. When bacterial cells increased and the QS system is fully activated, the biofilm matures and the production of ralstonins becomes high. Once *F. oxysporum* chlamydospores are formed, RSSC cells begin to detach from the biofilms, and some invade the spores. RSSC cells invade the chlamydospores and proliferate there. The infection kills the chlamydospores, and RSSC cells start migrating to the surrounding hyphae.

**Figure 9.**
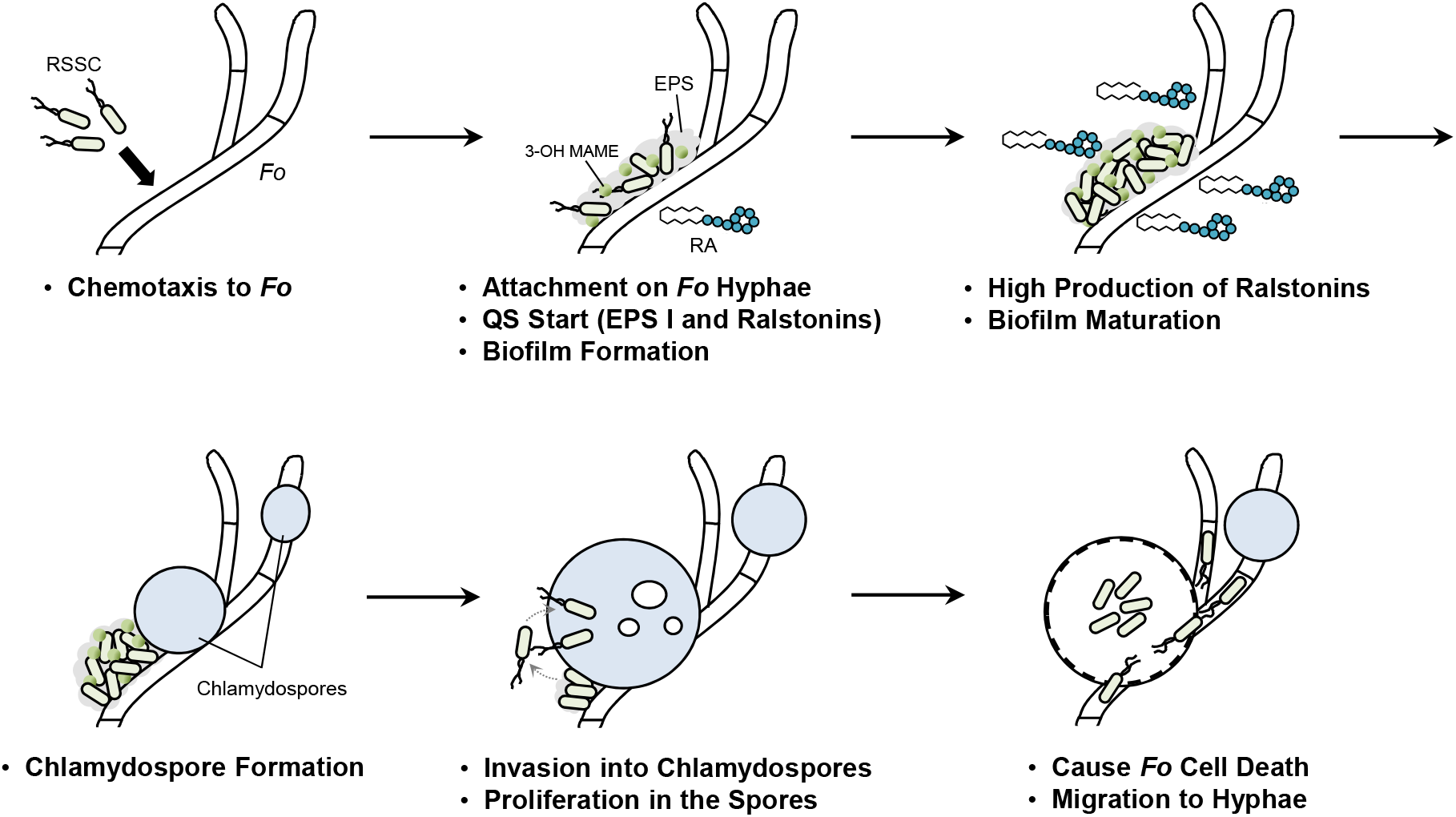
Proposed mechanism of RSSC endoparasitism in *F. oxysporum. Fo: F*.*oxysporum*, RA: ralstonin A.

The *phc* QS of RSSC strains is a critical system for parasitism in plants is properly activated at each step of infection, allowing them to invade and multiply within the hosts (18, 19). For example, the *phc* QS system regulates not only the production of the virulence factors (e.g., EPS I and plant-cell-wall-degrading enzymes) necessary for plant infection but also causes the metabolic changes to adapt the infection stages (37, 38). We found that in the RSSC endoparasitism on *F. oxysporum*, the *phc* QS system regulates biofilm formation on the hyphae and induces the chlamydospore formation of *F. oxysporum*. The production of ralstonins and EPS I, keys in these events, is under the QS control. Alternatively, T2SS and T3SS, also regulated by the *phc* QS system (37, 38), were not vital in this endoparasitism. This was different from the invasion strategy of *M. rhizoxinica* into the hyphae of *R. microsporus* (34). HrpG was reported to be partially involved in producing the QS-dependent factors (such as EPS I) (40), which may be the reason for the slightly reduced endoparasitism in Δ*hrpG*. Therefore, corporative cell–cell activities regulated by the *phc* QS system plays vital roles in this lethal interaction with the fungus.

Biofilm formation by bacteria on fungal mycelia has been occasionally reported (58–60). However, it was discovered that biofilm are essential for bacterial endoparasitism within fungi. Particularly, EPS I and ralstonins were found to be essential for RSSC to form biofilms on *F. oxysporum* hyphae. The overexpression of *xpsR* increased EPS I production and biofilm formation frequency. This resulted in a significant increase in the invasion rate of the recombinant compared with that of the wild-type strain. The EPS I around the bacterial cells was expected to increase the affinity of RSSC cells to *F. oxysporum* hyphae and wrap the aggregated cells there. The structure and amount of EPS I appear to be essential for the frequency and maturity of biofilms. Galactosaminogalactan was reported to contribute to the self-aggregation of *Aspergillus oryzae* hyphae (61, 62). A similar mechanism might be involved in increasing the affinity of EPS I for *F. oxysporum* hyphae. RSSC lipopeptides appeared to function as a biosurfactant and contribute to the biofilm formation. We cannot say for certain about the structural specificity needed for biofilm formation. Even mutants deficient in EPS I and/or lipopeptide production showed endoparasitism into *F. oxysporum* in the presence of the wild-type OE1-1. These factors are unlikely to be involved in the process after the biofilm formation.

Our analyses show that the produced RSSC lipopetides are an essential factor in determining whether the RSSC strain can be an endoparasite for *F. oxysporum*. The ralstonin-producing strains (OE1-1 and GMI1000) will likely be the best endoparasites among the tested strains. The ralstoamide-producing strains (MAFF 211479 and MAFF 211516) and the ralstopeptin-producing strain (MAFF 211519) are not parasitic on *F. oxysporum*. When ralstonins were supplied from OE1-1, the latter strains were also parasitized. A bioinformatic tool search did not find any bacteria (other than *Ralstonia*) producing a lipopeptide with a chemical structure similar to ralstonins. Fengycin A has chlamydospore-inducing activity against *F. oxysporum* and restores the parasitic potential of Δ*rmyA*, but the producer *Bacillus* spp. was not reported to invade *F. oxysporum* cells (54, 63). The discovery of further bacteria that can induce chlamydospores and endoparasitism against fungi may help to investigate whether the mechanism of RSSC’s endoparasitism is more ubiquitous.

Although our knowledge of the interaction between RSSC and *F. oxysporum* has deepened, how RSSC invades the *F. oxysporum* chlamydospores was not entirely clear in this study. According to our analysis, T2SS and T3SS are not involved in this endoparasitism. This was not our initial expectation. Some lipopeptides act as cell toxins by aggregating to form pores in plant cell membranes to facilitate nutrient exit or to disturb the ion balance inside in the cell (64–66). Although ralstonin A treatment-induced chlamydospore formation in *F. oxysporum*, such membrane disruption could not be seen from TEM and microscopic analyses. Instead, many mycelia changed to chlamydospores, and the cell membranes (or walls) became thicker. Thus, it is unknown the period when the cell membrane is disrupted. Endocytosis is an example of a mechanism of bacterial uptake by fungal cells (67). However, although several sections were observed by TEM, so far, no images reported the involvement of endocytosis. The possibility remains that we miss getting an image of the moment. To elucidate this mechanism, it will be important to examine microscopic observations at finer time intervals and the responses of *F. oxysporum* to the RSSC’s parasitism.

While this study was being prepared, a new paper on the parasitism of RSSC into a fungus was reported by Prof. Keller’s (63). There, the importance of chlamydospore induction by ralstonin A was shown using a different approach compared to this study. Additionally, several Gram-negative bacteria acted as a “hitchhiker” with RSSC cells and could invade the chlamydospores of the different host *Aspergillus flavus*. They reported that this event could be triggered simply by treating ralstonin A instead of RSSC cells. This is inconsistent with this study’s experimental results, in which *E. coli* failed to infect *F. oxysporum* by simultaneous inoculation with wild-type RSSC cells. Even RSSC cells cannot invade *F. oxysporum* chlamydospores if QS or related genes are deficient. It did not recover with ralsotonin alone. Likely, the difference in the hosts (*F. oxysporum* vs *A. flavus*) has a significant impact. Additionally, as confirmed in the *M. rhizoxinica*–*R. microsporus* system, the surface components of bacteria, such as LPS may be involved in host recognition (14). However, we have no idea on this respect in RSSC *F oxysporum* system. The conservation of the *phc* QS-dependent endoparasitism discovered here requires further verification in the future. The same is true regarding the “hitchhiker” behavior.

What advantages would RSSC obtain by parasitizing the fungus? RSSC strains must continue to survive in the soil even without a suitable plant host. There could be a situation where RSSC cells are present in the soil of a tomato field, but there are no tomato plants nearby to be infected. Fungi may be more ubiquitous, so the endoparasitism into fungi may be a good survival strategy for RSSC strains. It might be able to reach the host plants through fungal mycelia. As Keller *et al*. reported (63), endoparasitism in fungal hosts may confer resistance to environmental stresses on RSSC cells. Of course, the acquisition of stress tolerance is probably an essential aspect of this endoparasitism. However, given the possibility that RSSC cells assimilate nutrients to the point of killing *F. oxysporum* cells, such parasitism within fungi may be a survival strategy in the soil. The relationship may vary depending on the type of host. Since both *F. oxysporum* and RSSC infect tomato plants (68, 69), we are interested to see how this endoparasitism will affect the emergence of these diseases.

In terms of pest control, we need better to understand the behaviors of RSSC cells in the soil and suggest appropriate control methods. There might be multiple interactions that we do not yet know about, such as RSSC parasitism and symbiosis with other organisms, some of which kill the host and some of which keep the host alive. Through previous and current studies (22), we have reported that the *phc* QS system controls the phytopathogenic and endofungal parasitic behaviors of RSSC. Thus, the *phc* QS system should be an attractive control target. Suppose the QS system can be controlled by chemicals or microbes. In that case, it may be possible to prevent bacterial wilt disease before it develops, and even after that, the host plant’s recovery may be accomplished. We plan to examine the effects of QS inhibitors (such as PQIs) (30) on this endoparasitism in detail.

Conclusively, we found a QS-dependent invasion of RSSC into *F. oxysporum*. We not only have confirmed the importance of ralstonins as *F. oxysporum* chlamydospore inducers but have also confirmed that EPS I production and biofilm formation are cleverly regulated by the *phc* QS system during the development of endoparasitism on *F. oxysporum* chlamydospores. The RSSC infection is likely a means of killing the *F. oxysporum* spores and obtaining nutrition. Consequently, RSSC appears to allow free movement within *F. oxysporum*. RSSC strains may be the phytopathogenic bacteria that have evolved to fit field conditions more cleverly than we imagine.

## Methods

### Nylon Net Sandwich Assay

GFP-expressing RSSC strains were inoculated into 2 × BG media (2 mL) containing kanamycin (50 μg/mL) and cultured overnight by shaking. A nylon net filter (11 μm, 47 mm, Merck) was placed on the surface of a BG agar medium (25 mL in 90 mm plastic dish) containing kanamycin (100 μg/mL). After the B medium (50 μL) was added to the top of the net, another nylon net (20 μm, 47 mm, Merck) was placed on top. RSSC cell suspension (5 μL) and *F. oxysporum* spore suspension (5 μL) were spotted on the top net, 15 mm apart from each other. The plate was incubated at 30°C for three days. The top net filter was removed from the medium and washed thoroughly with MilliQ water to remove the bacteria cells. Chlamydospores were collected by rubbing the surface using a cover glass. The collected chlamydospores were washed with MilliQ water (0.5 mL) and observed under a fluorescence microscope (BZ-X810, Keyence). The invasion rate per condition was calculated from the three to five dishes.

### Biofilm Formation Assay

RSSC strains were incubated in CPG media (2 mL) at 30°C overnight. The cultures were transferred to 2 mL tubes and centrifuged at 15,000 ×*g* for 3 min. The old media were discarded and the bacterial cells were suspended in CPG media (1 mL), and the OD_600_ was adjusted to 0.1. New CPG medium (95 μL) and bacterial cell suspension (5 μL) were dispensed into each well of a 96-well microtiter plate (Thermo Fisher Scientific). The plates were incubated at 30°C for 20 h. To the culture medium, 1% (w/v) crystal violet solution (25 μL) was added, allowed to stand at room temperature for 25 min, and then excess crystal violet was removed with a pipette. The well was then washed twice with MilliQ water (200 μL). Ethanol (200 μL) was added and allowed to stand for 10 min to extract the crystal violet. The solution was transferred to a flat-bottomed 96-well plate (Iwaki), and the absorbance at a wavelength of 595 nm was measured using a plate reader (Multiskan FC, Thermo Fisher Scientific).

### Glass-Bottom Dish Assay

Hot BG agar medium (600 μL, 1% (w/v) agar) is dispensed into a glass-bottom dish (Matsunami). After cooling, *F. oxysporum* spore suspension (2 μL) was spotted on the surface of BG agar. The dish was incubated for 36 h at 30°C. The RSSC cell suspension in BG medium (OD_600_= 0.25, 2 mL) was pipetted onto the surface of the *F. oxysporum* colony in the dish. The BG medium was removed by pipetting after incubation for 9 or 24 h at 30°C. By washing with by washing with MilliQ water (1 mL) twice, the excess RSSC cells were discarded. If necessary, a staining operation was added before washing. DBA-FITC (VECTOR LABORATORIES) and SYTO9 (Thermo Fisher Scientific) were used to stain EPS I and bacterial cells, respectively. The dish was observed using a fluorescence-inverted microscope (BZ-X810).

### Construction of Gene-Deletion Mutants

A synthetic gene was prepared by fusing about 500 bp upstream and about 500 bp downstream of the target gene (Eurofins Genomics). The fused gene was released from the provided vector by restriction enzyme digestion and introduced into the pK18 mobsacB vector. This plasmid was electroporated into RSSC cells (OD_600_= 0.6, in 10% glycerol), and the cells were incubated in SOC medium (1 mL, Sigma-Aldrich) for 6 h at 30°C. Also, kanamycin-resistant (50 μg/mL) and sucrose-sensitive (10%) recombinants were selected. The recombinants were incubated in SOC medium for 4 h, with a kanamycin-sensitive (50 μg/mL) and sucrose-resistant (10%) recombinant. Furthermore, the deletion of the target region was confirmed by PCR. The other mutants were previously reported by our group or were gifts from Prof. Hikichi (Kochi University).

### Isolation of Ralstonin A

OE1-1 cells grown in B medium for six hours was spread on 1000 BG agar plates, and the plates were incubated at 30°C for three days. BG agar was cut into small pieces and soaked in acetone (20 L) for two hours. The acetone extract was collected by filtration, and the agar residue was soaked again in acetone (20 L) for two hours. The combined extracts were evaporated to remove acetone, extracted thrice with EtOAc, and dried over Na_2_SO_4_. The EtOAc extracts were concentrated and chromatographed on a silica gel column, and eluted stepwise with solvents of increasing polarity from *n*-hexane to EtOAc, and then from EtOAc to MeOH. The fractions containing ralstonins (60% and 100% MeOH eluates) were subjected to HPLC with an InertSustain C18 column (250 mm × 10 mm, 5 μm) at a flow rate of 4 mL/min with 45% MeCN in 0.1% aq. trifluoroacetic acid (TFA) to yield ralstonin A (13 mg).

### TEM Observation

*F. oxysporum* chlamydospores induced by ralstonin A or by coculturing with strain OE1-1 were used. Ultrastructural analyses of the chlamydospores were conducted by Tokai-EMA (Japan).

### Statistical Analysis

Mean values and standard errors (SEM) were calculated using GraphPad Prism 6.0. Levels of significance were evaluated using one-way ANOVA, followed by Tukey’s multiple comparison tests by GraphPad Prism 6.0.

## Acknowledgments

We thank Prof. Y. Hikichi (Kochi University) for his valuable advice and comments. This study was supported by JSPS KAKENHI Grants No. 17K19244, 22H02275, and 22H04888.

## Figure and Legends

**Figure S1.**
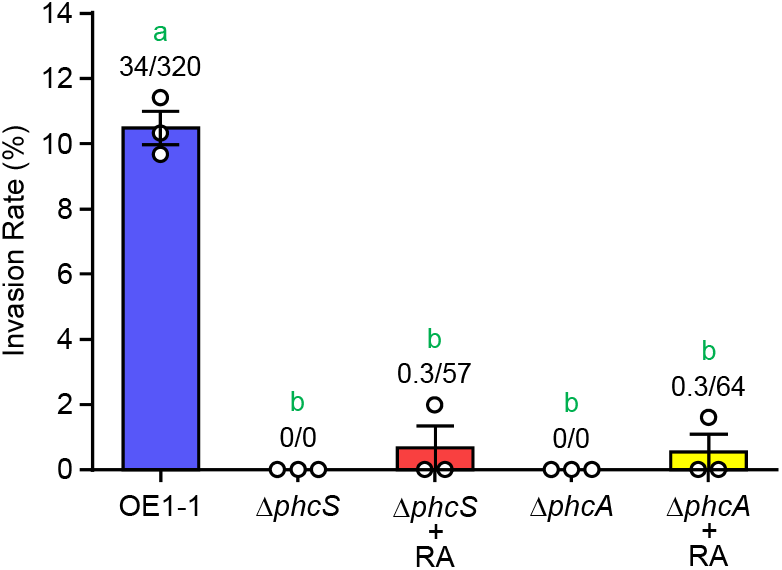
Invasion rates of Δ*phcS* and Δ*phcA* and their response to ralstonin A. Ralstonin A (RA) was used at 0.1 nmol/disc. OE1-1 is a positive control. The error bars show the mean ± SEM (*n* = 3). The numbers above the bars indicate infected spores/total spores. The green letters above the bars show significant differences (*p* < 0.05, Tukey’s test).

**Figure S2.**
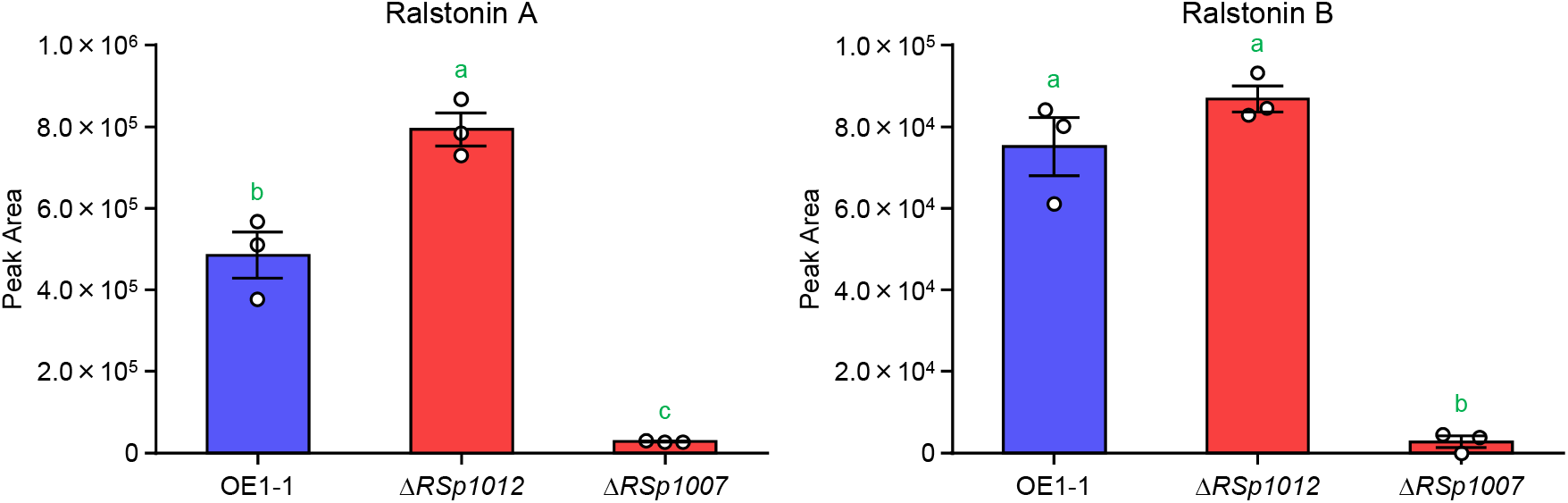
Comparison of ralstonin production in OE1-1, Δ*RSp1012*, and Δ*RSp1007*. On the left are the data for Ralstonin A and on the right are the data for Ralstonin B. The error bars show the mean ± SEM (*n* = 3). The green letters above the bars show significant differences (*p* < 0.05, Tukey’s test).

**Figure S3.**
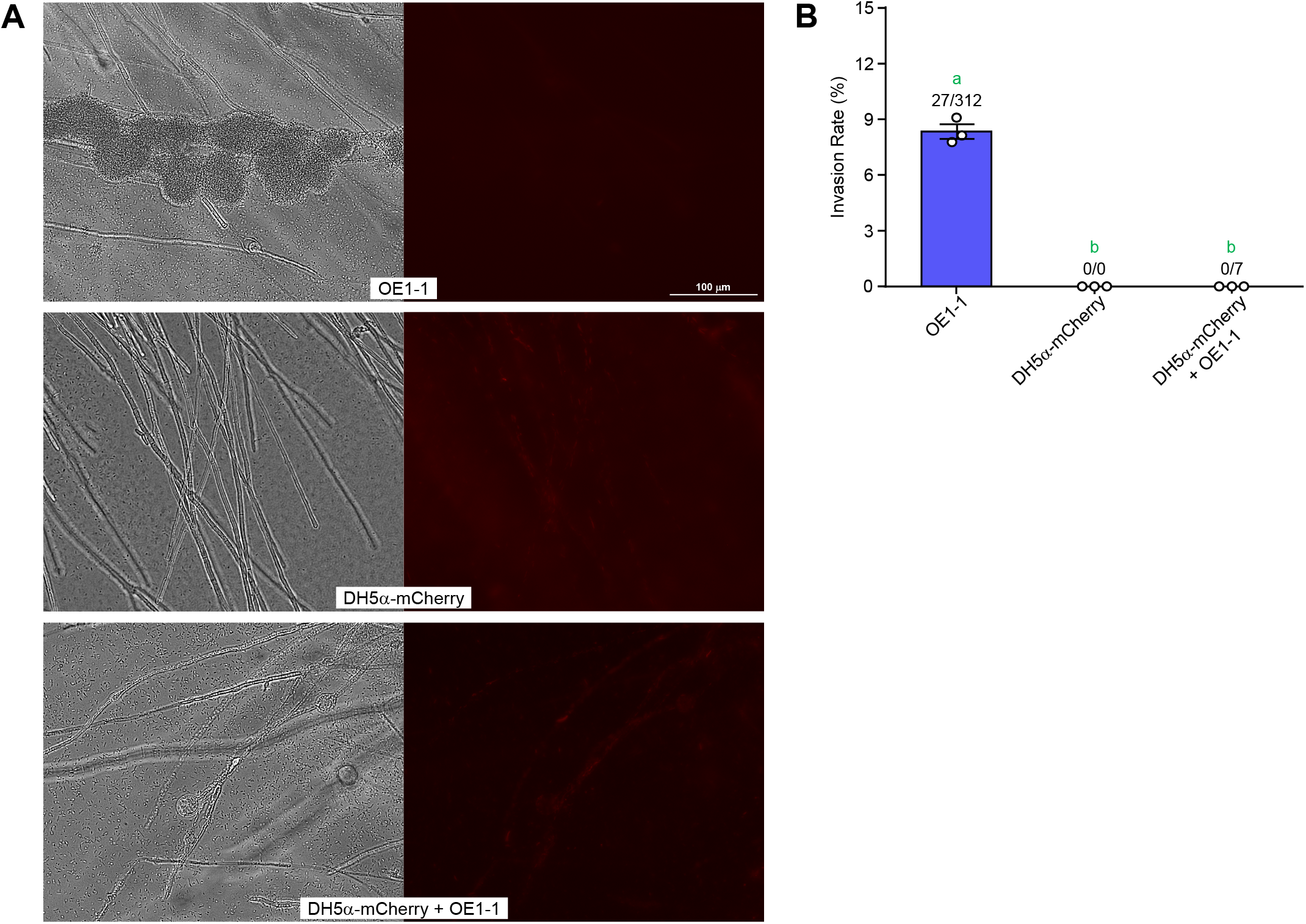
Evaluation of biofilm formation and invasion abilities of *E. coli* DH5α-mCherry. (A) The photos of coculturing of *F. oxysporum* with OE1-1, *E. coli* DH5α-mCherry, and both. The bright field image is on the left, and the fluorescence image is on the right. (B) Invasion rates of OE1-1 and *E. coli* DH5α-mCherry into *F. oxysporum* chlamydospores. *E. coli* DH5α-mCherry was not parasitized even when cocultured with OE1-1. The error bars show the mean ± SEM (*n* = 3). The numbers above the bars indicate infected spores/total spores. The green letters above the bars show significant differences (*p* < 0.05, Tukey’s test).

**Figure S4.**
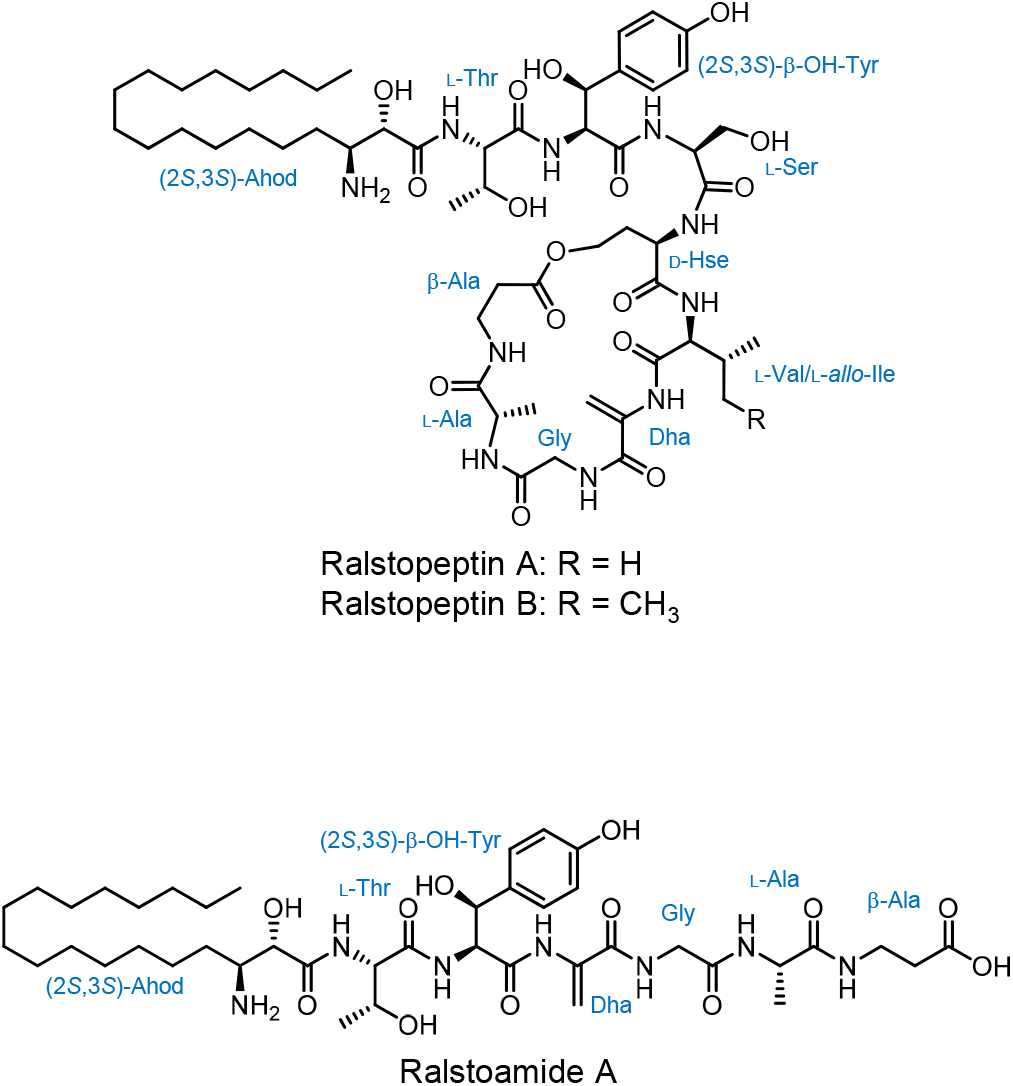
Structures of ralstopeptins A, B, and ralstoamide A.

